# R-loop-induced p21 expression following CDC73, CTR9, and PAF1 loss protects cancer cells against replicative catastrophe following WEE1 inhibition

**DOI:** 10.1101/2021.07.14.452205

**Authors:** Linda van Bijsterveldt, Helga B. Landsverk, Viola Nähse, Samuel C. Durley, Sovan S. Sarkar, Randi G. Syljuåsen, Timothy C. Humphrey

## Abstract

WEE1 inhibitors have now advanced into clinical studies as monotherapy or in combination with chemoradiotherapy in *TP53, RAS, BRAF*, and *SETD2* mutation carriers across several tumour types, yet mechanisms of resistance are still poorly understood. Here, we further elucidate the mechanisms by which AZD1775, the most potent WEE1 inhibitor, kills cells and reveal additional genetic interactions that can result in resistance, but could be used to optimise its clinical utility.

We identified RNA Polymerase II-associated factor 1 (PAF1) complex members, CDC73, CTR9, and PAF1 as major determinants of WEE1-inhibitor sensitivity in isogenic *SETD2*-positive and negative cell lines. *PAF1*-knockdown cells resist higher doses of the WEE1 inhibitor, which we show is due to reduced DNA damage induction (γH2AX) and delayed G1 checkpoint activation, ultimately protecting cells against replicative catastrophe. Investigations into the molecular mechanisms responsible for PAF1-mediated resistance identify involvement of R-loops and subsequent activation of the cyclin-dependent kinase inhibitor p21Cip1/Waf1, which in addition to causing prolonged G1 arrest in the following cell cycle, also regulates CDK activity, therefore limiting replication.

These results provide evidence that the PAF1 complex and p21 are important regulators of proliferation under increased DNA replication stress and their expression levels might be useful biomarkers to predict clinical response to WEE1 inhibitors and other ribonucleotide reductase inhibitors.

## Introduction

The use of cell cycle checkpoint kinase inhibitors, including WEE1 inhibitors, has emerged as a promising targeted strategy aimed to exploit cancer cells with a dysfunctional G1/S checkpoint (van Bijsterveldt *et al*., 2020). Several WEE1 inhibitors have been developed pre-clinically (Matheson, Backos and Reigan, 2016), and a number of those have now advanced into clinical studies as monotherapy or in combination with chemoradiotherapy in *TP53, RAS*, and *BRAF* mutation carriers across several tumour types (Rorà *et al*., 2020). WEE1 kinase is responsible for inhibitory tyrosine-15 (Y15) phosphorylation of the ATP-binding domain of cyclin dependent kinase 1 and 2 (CDK1/2) (Parker and Piwnica-Worms, 1992). Accordingly, inhibition of WEE1 results in CDK1 and CDK2 hyperactivation during S-phase, which promotes increased replication origin firing as well as degradation of RRM2, the regulatory subunit of ribonucleotide reductase (RNR) that catalyses the rate-limiting step of the formation of deoxyribonucleotides (dNTPs) (D’Angiolella *et al*., 2012; Pfister *et al*., 2015).

Our group has recently demonstrated that *SETD2*-deficient cancers, with SETD2 encoding the only histone methyltransferase that catalyses H3 lysine 36 trimethylation (H3K36me3) (Edmunds, Mahadevan and Clayton, 2008), can be targeted by exploiting a synthetic lethal interaction with WEE1 inhibition (WEE1i) by AZD1775 (Pfister *et al*., 2015). *SETD2*-deficient cells display lower RRM2 transcript levels and can therefore be preferentially targeted with the potent WEE1 inhibitor AZD1775 as dNTP pools reach critically low levels (Pfister *et al*., 2015). Reduced intracellular dNTP pools in this context provoke catastrophic widespread fork collapse and accumulation of DNA double-strand breaks (DSBs), ultimately leading to replicative catastrophe (Poli *et al*., 2012; Pfister *et al*., 2015). Therefore, cell death resulting from WEE1 inhibition in combination with the loss of SETD2 is thought to be primarily caused by replicative catastrophe (Pfister *et al*., 2015; Pai *et al*., 2019). Even though synthetic lethality-based cancer therapies have been proven to be highly effective, therapy outcomes may not be durable due to recurrent patterns of resistance development within tumours. In the case of WEE1 inhibition, it remains unclear whether processes beyond canonical checkpoint control can modulate response and resistance to WEE1 inhibitor treatment, which is why there is a pressing need to identify clinically relevant biomarkers of sensitivity.

The RNAPII-associated factor 1 (hPAF1) complex, is a positive elongation factor and is also required for chromatin modifications, particularly ubiquitination of H2B K120 and H3 K4 methylation (Nevan J. Krogan *et al*., 2003; Ng *et al*., 2003; Wood *et al*., 2003). In *S. cerevisiae* and *S. pombe* but not mammalian cells, recruitment of Set2 and H3K36 methylation to actively transcribed genes relies on components of the yeast Paf1 complex, as well as the RNAPII CTD kinase Ctk1 (ortholog of CDK12) in *S. cerevisiae* and cyclin-dependent kinase 9 (Cdk9) in *S. pombe* (Nevan J Krogan *et al*., 2003; Mbogning *et al*., 2013). Our group has identified *PAF1* and *PRF1* (assigned RTF1 in mammalian cells) (Vos *et al*., 2020), two components of PAF1C, as two genes putatively responsible for suppression of the *set2*Δ *wee1-50* synthetic lethality using a suppressor screen. The human PAF1C mainly functions to release paused RNAPII from the promoter-proximal pause site (20-60 nucleotides downstream of the transcription start site (TSS)) into effective transcriptional elongation as it displaces negative elongation factor (NELF) from the RNAPII funnel (Chen *et al*., 2015; Yu *et al*., 2015; Vos *et al*., 2018). Inactivation of the PAF1 complex leads to RNAPII accumulation at the proximal-promoter pausing site in both budding yeast (Wahba *et al*., 2011) and mammalian cells, causing unscheduled R-loop formation (Shivji *et al*., 2018) and checkpoint activation. To prevent the genome against R-loop associated DNA damage, unscheduled R-loop accumulation can activate the ATR-Chk1 signalling pathway in a replication-dependent manner (Matos *et al*., 2020), thereby enforcing checkpoint arrest, suppression of transcription-replication conflicts, and promoting fork recovery. Dysregulation of R-loops can cause genomic instability, one of the enabling characteristics of cancer, via formation of DSBs, chromosome breaks, and recombination events (Hanahan and Weinberg, 2011; Crossley, Bocek and Cimprich, 2019).

Here, we find that loss of *paf1* in fission yeast suppresses the *set2*Δ *wee1-50* synthetic lethality. Furthermore, depletion of PAF1C members, including CDC73, CTR9 and PAF1 in human osteosarcoma (U2OS) cells results in resistance to cell death following WEE1 inhibition in both the presence and absence of SETD2. Depletion of PAF1C members results in an increase in p21, which binds to and inhibits both cyclin-CDK complexes and proliferating nuclear antigen (PCNA). The upregulation of p21, via its ability to induce the G1 checkpoint following DNA damage, prevents replication and protects cells from cell death following WEE1 inhibition. These results are of importance to guide the future clinical implementation of the WEE1 inhibitor, as tumours with low p21 expression as a result of transcriptional dysregulation or *TP53* deficiency may be particularly sensitive, whereas tumours with mutations in *CTR9, PAF1, CDC73*, or other cancers with mutations up-regulating p21 levels might prove to be resistant to ribonucleotide reductase inhibitor treatment.

## Results

### Loss of transcription elongation factor Paf1 rescues the synthetic lethality resulting from Set2 deletion and Wee1 inactivation in *S. pombe*

To explore the possible mechanisms of resistance to WEE1 inhibitor treatment in H3K36me3-deficient cells, a genetic screen was performed in fission yeast to isolate spontaneous mutants that suppressed the synthetic lethal interaction between *set2Δ* and *wee1-50*, a temperature-sensitive *wee1*, which is inactive when grown at the restrictive temperature of 35.5°C (Russell and Nurse, 1987). Of approximately 10,000 *set2Δ wee1-50* cells plated, 23 mutants were identified which were able to grow at 35.5°C. Sequence analysis of these mutants included loss of function mutations in *cdc2*, as expected, and a number of additional mutants, including mutations in seven genes involved in transcription, one of which was *paf1*, a subunit of the polymerase associated factor complex.

To validate the results of the screen, we generated *set2Δ wee1-50 paf1Δ* and tested their ability to form colonies using a serial dilution assay (Figure 1A). Growth of *set2*Δ *wee1-50* was compromised when grown at semi-permissive (32°C) temperature (Figure 1A) as previously described (Pai *et al*., 2019). Deletion of *paf1*^*+*^, however, suppressed the *set2Δ wee1-50* synthetic lethality, as the *set2Δ wee1-50 paf1Δ* triple mutants were able to grow at 32 °C in contrast to *set2Δ wee1-50* double mutants (Figure 1A).

**Figure 1.**
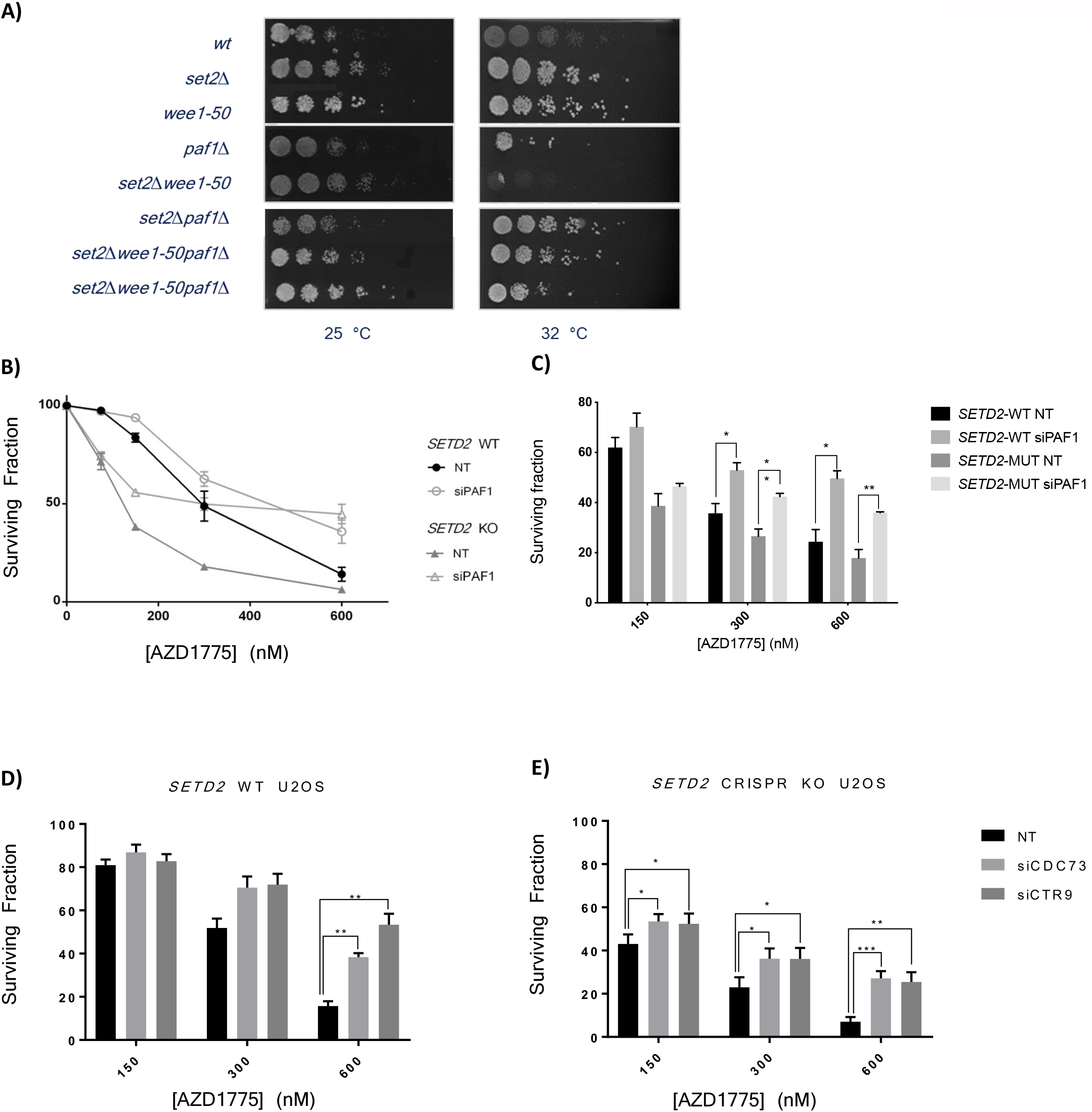
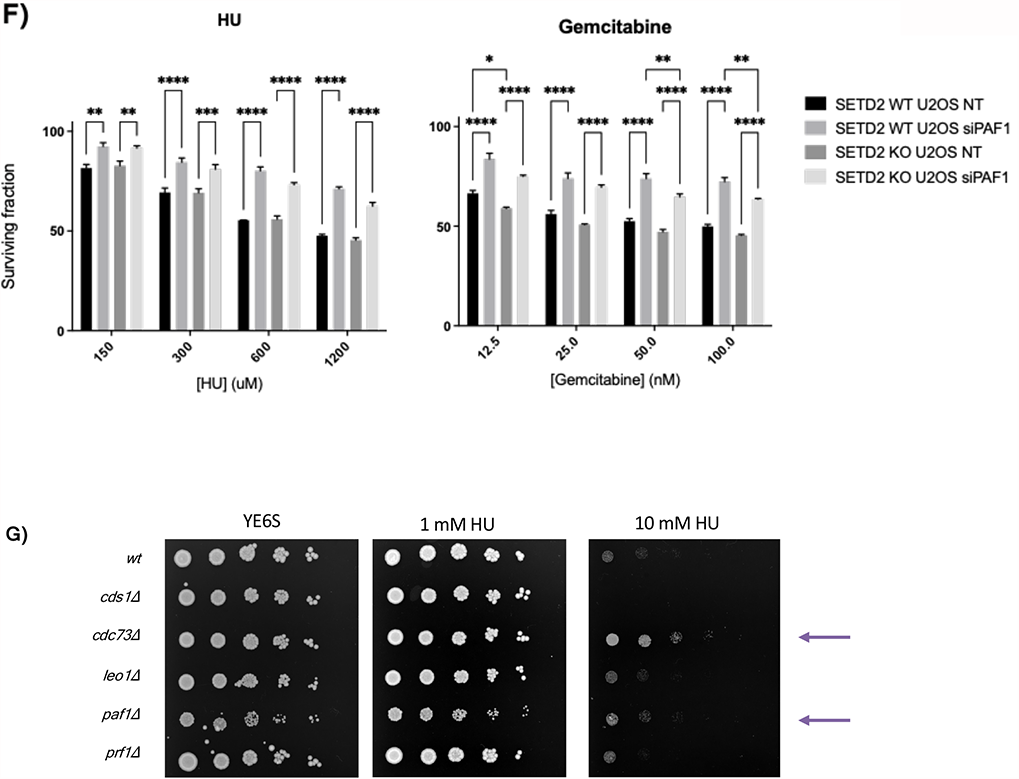
PAF1C-mediated resistance to depleted dNTP pools. **(A)** The growth response of wild-type *S. pombe*, set2Δ, wee1-50, paf1Δ, set2Δ wee1-50, set2Δ paf1Δ, and set2Δ wee1-50 paf1Δ was compared by spotting 5-fold serial dilutions of each strain onto YE6S medium. Plates were the incubated for an appropriate time at either 25, 32 or 35.5 °C. (**B-C**) *SETD2* WT and *SETD2* CRISPR KO U2OS cells after 72 h exposure to different concentrations of AZD1775 (75, 150, 300 and 600 nM). **(D-E)** Viability assay to examine the impact of other PAF1 complex factors (including CDC73 and CTR9) silencing on AZD1775-sensitivity. Survival fraction of non-targeting control, CDC73, and CTR9 siRNA treated *SETD2* WT and *SETD2* CRISPR KO cells after 72 h exposure to different concentrations of AZD1775 (75, 150, 300 and 600 nM). Data points and bars represent the mean and SEM of ≥ three independent experiments; * P < 0.05; ** P < 0.01; and *** P < 0.001. **(F)** Viability assay to examine the impact of PAF1 silencing on HU or gemcitabine. Survival fraction of non-targeting control and PAF1 siRNA treated *SETD2* WT and *SETD2* CRISPR KO U2OS cells after 72 h exposure to different concentrations of HU (150, 300, 600 and 1200 uM) or gemcitabine (12.5, 25, 50 or 100 nM). Data points and bars represent the mean and SEM of ≥ three independent experiments; * P < 0.05; ** P < 0.01; and *** P < 0.001. **(G)** Viability assay to examine the sensitivity to HU in gene deletions of PAF1C components in S. *pombe*.

### Knockdown of PAF1, CTR9, or CDC73 desensitises U20S cells to WEE1-inhibition

To investigate whether Paf1-mediated suppression of Wee1 inactivation is conserved from fission yeast to human cells, PAF1 was silenced by RNA interference in *SETD2* WT and *SETD2* CRISPR knockout (KO) human osteosarcoma (U20S) cell lines and sensitivity to the WEE1-inhibitor AZD1775 was measured. Knockdown of PAF1 was confirmed by Western Blot (Supplemental Figure 1A). Knock-down of PAF1 significantly restored cell viability in the presence 150, 300 and 600 nM of AZD1775 (Figure 1B-C). The survival rate at maximum inhibition (Imax% value) differed significantly between non-targeting control siRNA and PAF1 siRNA treated U2OS cells, with a stronger effect observed in *SETD2* CRISPR KO U2OS cells (non-targeting siRNA control I_max%_ = 6.36 versus PAF1 siRNA I_max%_ = 44.7, p-value= 0.0082) (Figure 1B-C).

To exclude the possibility that U20S cells have acquired other unaccounted mutations that may contribute to the resistant phenotype observed following PAF1 knock-down, we also silenced PAF1 in the renal cell carcinoma (RCC) cell lines 786-0 and A498 (bearing a truncation mutation in *SETD2*), where WEE1-inhibitor sensitivity was measured (Supplemental Figure 1B). Consistent with our previous results, knock-down of PAF1 significantly desensitised *786-0* and *SETD2-*mutant cell line A498 to AZD1775 (Supplemental Figure 1B). Increased cell viability following AZD1775 treatment in siPAF1-treated *SETD2* CRISPR KO U2OS was, however, not accompanied by caspase-3-mediated-apoptosis resistance (Supplemental Figure 1C). The increased cell viability following PAF1 KD is therefore not likely to result from resistance to cell death induction by apoptosis.

To further characterise the role of the PAF1 complex in mediating resistance to WEE1 inhibition, siRNA mediated gene-silencing was used to assess the effect of depletion of individual PAF1 complex members on AZD1775-sensitivity in both *SETD2* WT and *SETD2* KO U2OS cells. Knock-down of CDC73 and CTR9 phenocopied the effect of PAF1 knockdown (Figure 1D-E) following WEE1-inhibitor treatment, whereas silencing of LEO1 and RTF1 did not have a significant effect on cell viability (data not shown). Knockdown of PAF1 complex components was validated using Western Blot (Supplemental Figure 1D). We can therefore conclude that WEE1-inhibitor sensitivity is dependent on the presence of at least a subset of PAF1 complex members, including CDC73, CTR9, and PAF1.

Moreover, to investigate the importance of the PAF1C in regulating the cellular response to replication stress, we analysed the sensitivity patterns to hydroxyurea (HU) and gemcitabine following PAF1 depletion in *SETD2* WT and *SETD2* CRISPR KO U2OS cells. Our results indicate that PAF1 depletion renders U2OS cells less sensitive to increasing concentrations of hydroxyurea (HU) and gemcitabine (Figure 1F), all of which directly deplete dNTP pools and subsequently block replication fork progression (Alvino et al., 2007; Cerqueira, Fernandes, & Ramos, 2007). Similarly, in *S. pombe*, a spot assay showed that *paf1Δ and cdc73Δ* display reduced sensitivity to high concentrations of HU when compared to a wild-type strain (Figure 1G). We therefore conclude that loss of PAF1C components confers an evolutionarily conserved cross-resistance among inhibitors of ribonucleotide reductase.

### Loss of PAF1 rescues AZD1775-induced replicative catastrophe through damage reduction

As previously described, AZD1775 leads to selective killing of SETD2 KO U20S cells through dNTP pool depletion, accumulation of non-replicating S-phase cells (exhibiting a DNA content between 2N and 4N, but not incorporating the synthetic nucleoside bromodeoxyuridine [BrdU]) and, ultimately leading to replicative catastrophe (Pfister et al., 2015). Therefore, we tested whether loss of PAF1 restored cell viability by alleviating AZD1775-induced S-phase effects, such as reduced BrdU incorporation, indicative of slow S-phase progression, and DNA damage.

To examine the progression through S-phase following PAF1 silencing in *SETD2* WT and *SETD2* CRISPR KO U2OS, cells were treated with 300 nM AZD1775 for 48 hours, pulse-labelled with BrdU for 30 min and collected for cell cycle analysis. AZD1775 treatment resulted in accumulation of 50 ± 0.3 % of the *SETD2* CRISPR KO U2OS cells in non-cycling S-phase, with only 16% in G1 (Figure 2A-B). In siPAF1-treated *SETD2* CRISPR KO U2OS cells, despite inhibition of WEE1, only 26.4 ± 3.8 % of cells were in non-replicating S-phase, whereas the fraction of cells in G1 phase of the cell cycle was increased (38.6 ± 2.2 %) (Figure 2A-B).

**Figure 2.**
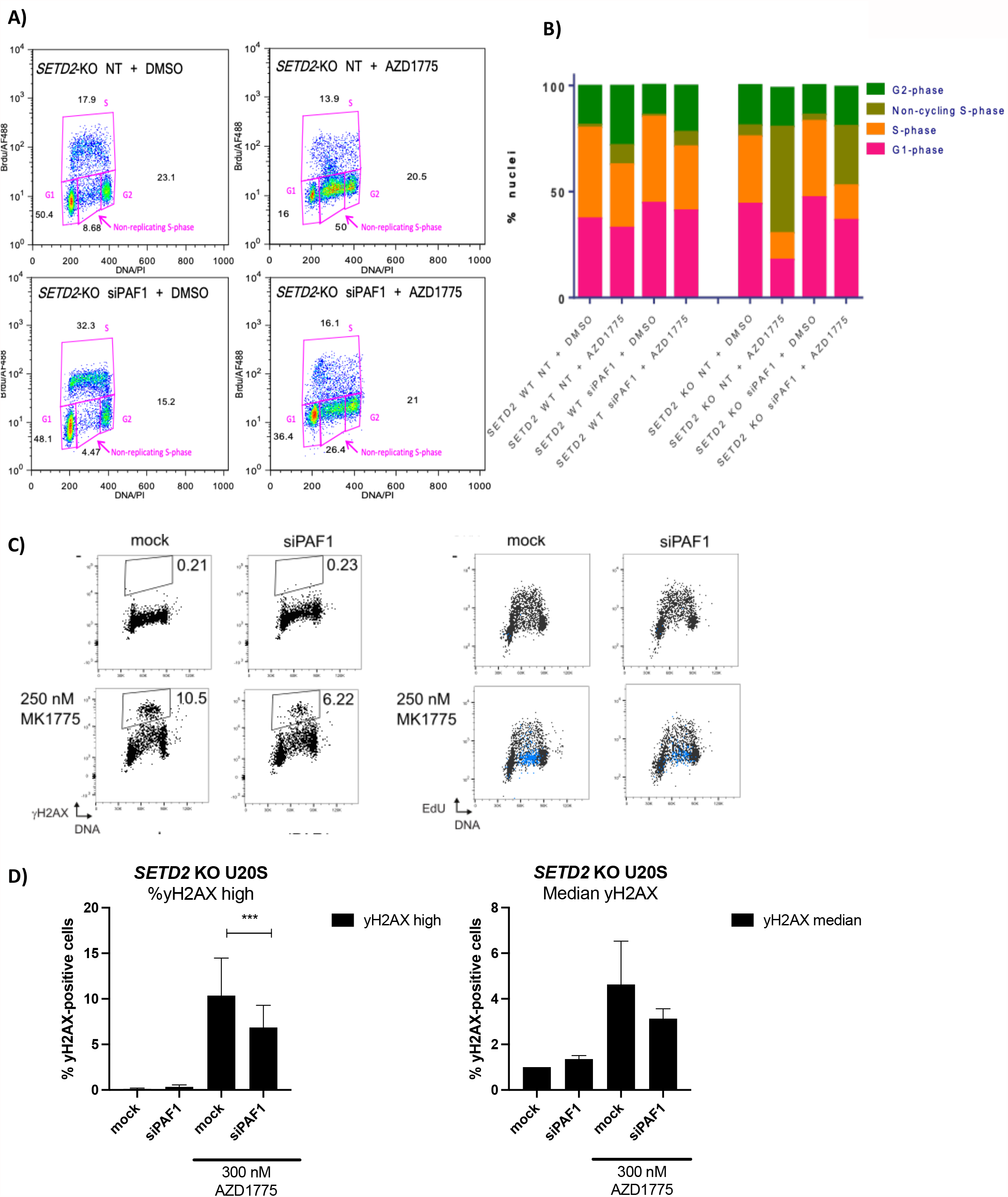
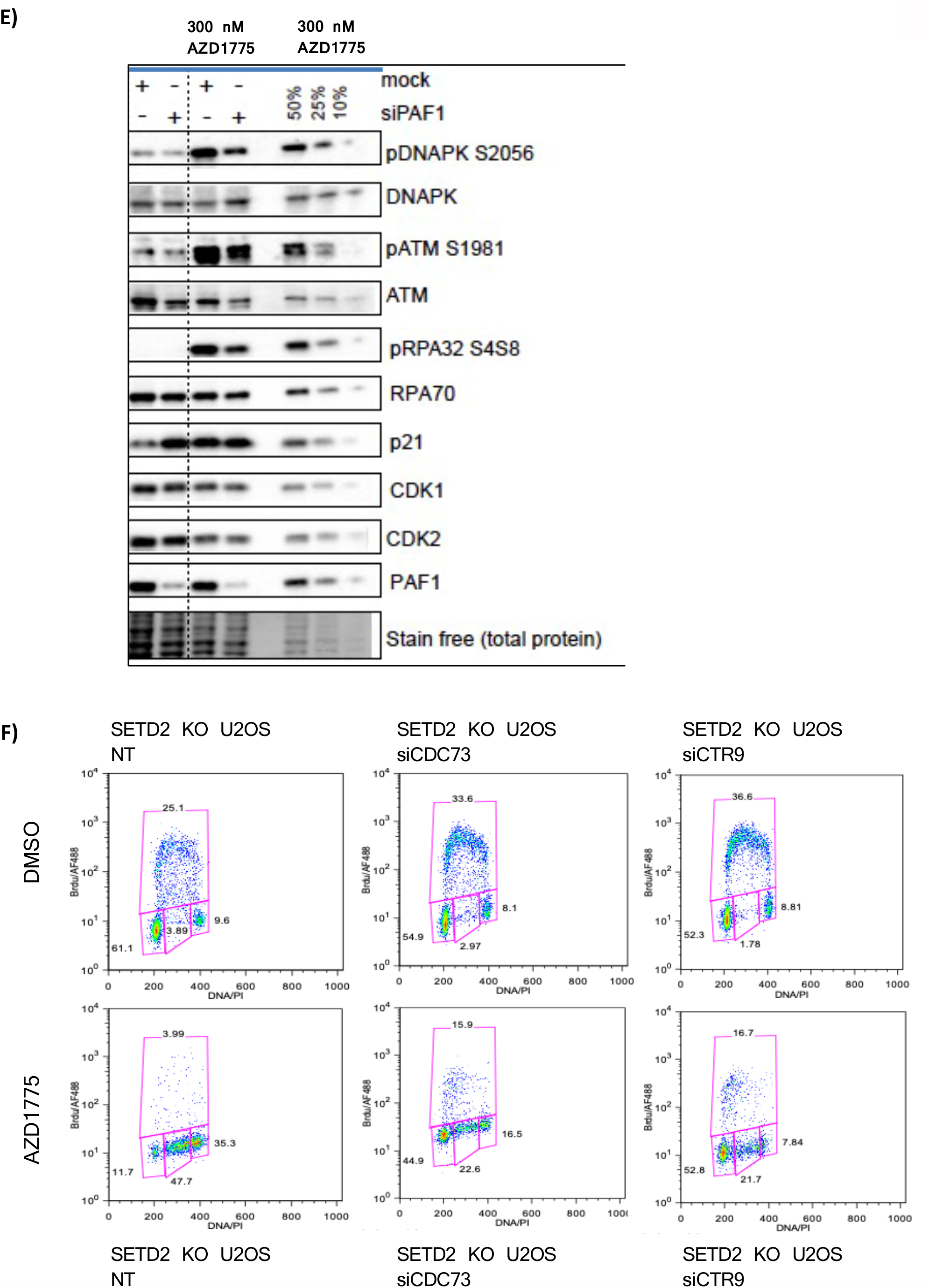
Loss of PAF1 results in G1 accumulation and decreased S-phase associated DNA damage. **(A)** 48 hours following DMSO or AZD1775-treatment (300 nM), control or siPAF1 treated *SETD2* KO U2OS cells were pulse-labelled with BrdU for 30 min and collected for cell cycle analysis. The data shown are from a single representative experiment out of three repeats. Numbers are the relative percentage of cell cycle stage. **(B)** Quantification of data shown in (A). **(C)** *SETD2 CRISPR KO* U2OS were treated with control or siPAF1 siRNA and exposed to 250 nM of AZD1775 for 48 hours. One representative experiment showing yH2AX vs DNA (left panel) and **EdU** vs DNA (right panel). The numbers indicate the percentages of cells in the high yH2AX-gated population (shown in black box). These cells correspond with the non-replicating S-phase cells as shown in the EdU plots (blue population). **(D)** Quantification of the yH2AX medium and high populations shown in (C). **(E)** Western blot analysis of *SETD2* CRISPR KO U2OS cells exposed to either non-targeting control siRNA or PAF1 siRNA and/or treated with either DMSO or 300 nM AZD1775. Lamin B was used as a loading control. **(F)** 48 hr following AZD1775-treatment, control, siCDC73, or siCTR9 treated *SETD2* KO U2OS cells were pulse-labelled with BrdU for 30 min and collected for cell cycle analysis.

To uncover the molecular mechanisms by which PAF1 depletion reduces the toxic effects of WEE1i in *SETD2* KO U20S cells, we assessed DNA damage, using an antibody to γH2AX, following WEE1 inhibitor treatment in S-phase U20S cells. Firstly, we found that replicative catastrophe resulting from AZD1775-treatment is a direct function of DNA damage. AZD1775 treatment (1 μM) resulted in the accumulation of two distinct populations, a high (16.1%) and an intermediate (21.7%) γH2AX population as early as 6 hours post-treatment (Supplemental Figure 2A). FACS analysis of DNA damage 4 hours following WEE1 inhibitor wash-out showed a clear reduction in the intermediate γH2AX population (reduction of 15.5%), but no change in the percentage of cells with the highest levels of damage (Supplemental Figure 2B), suggesting that some of the DNA damage breaks might be irreversible, or take longer to repair. When the high γH2AX population was gated (indicated in black in the EdU plot, Supplemental Figure 2A), we found that the high γH2AX population was EdU-negative, suggestive of replication arrest (Supplemental Figure 2B).

To test whether the alleviation of replicative catastrophe following PAF1 depletion was a direct result of reduced DNA damage (γH2AX) in S-phase, we compared γH2AX levels 48 hours following AZD1775-treatment (250 nM) in siPAF1 treated U20S with control siRNA treated cells. We found that depleting PAF1 resulted in significantly reduced levels (p = 0.0038) of γH2AX-high cells (6.85%) compared to siNT-treated *SETD2* KO U20S cells (10.36%) following addition of the WEE1-inhibitor AZD1775 (Figure 2C-D). Furthermore, as shown in Figure 2E (quantified in Supplemental Figure 2C), we found that levels of phosphorylated ATM, DNA-PK and RPA32 were also reduced following AZD1775 treatment in *SETD2* KO U20S cells treated with siPAF1 when compared to the control-treated cells. This suggests reduced or delayed formation of replication-associated DNA DSBs (Liaw, Lee and Myung, 2011) and altered damage signalling in the absence of PAF1.

Finally, we assessed the effect of depleting other PAF1C components on the non-replicating S-phase population following WEE1 inhibition and found that depletion and CDC73 and CTR9 resulted in a similar reduction in the non-replicating S-phase population. When U20S cells depleted for CDC73 or CTR9 were treated with AZD1775 for 48 hours, we also observed an accumulation of cells in G1 and a decrease in S-phase population, with an effect on the rate of DNA synthesis with lower BrdU incorporation (22.6% and 21.7% BrdU-negative S-phase cells for CDC73 and CTR9, respectively), when compared to the control-siRNA treated *SETD2 CRISPR KO* U20S cells (47.7%) (Figure 2F).

Taken together, these results show that depletion of CDC73, CTR9 or PAF1 result in a larger fraction of cells in G1 phase following AZD1775 treatment, as well as a reduction in the non-replicating S-phase fraction, due to a lower percentage of cells with median and high γH2AX levels, leading to an increase in cell viability in the context of WEE1 inhibition.

### R-loop and P53-dependent up-regulation of CDKN1A following loss of CDC73, CTR9, and PAF1

To further investigate the proliferation defect observed after CDC73, CTR9, or PAF1 depletion, siRNA-transfected U2OS cells were synchronised at the G1/S border by double thymidine block followed by release and assessment of cell cycle progression into S-phase by flow cytometry (Supplemental Figure 3A-B). Consistent with the growth assays (Supplemental Figure 2), FACS analysis indicated that *SETD2* WT U2OS cells depleted of PAF1, CDC73 or CTR9 spent more time in G1 after release than siNT-treated control cells (Figure 3A-B, Supplemental Figure 3A-B). These data prompted us to examine the relationship between p21, a well-studied regulator of the G1/S transition, and components of the PAF1C.

**Figure 3.**
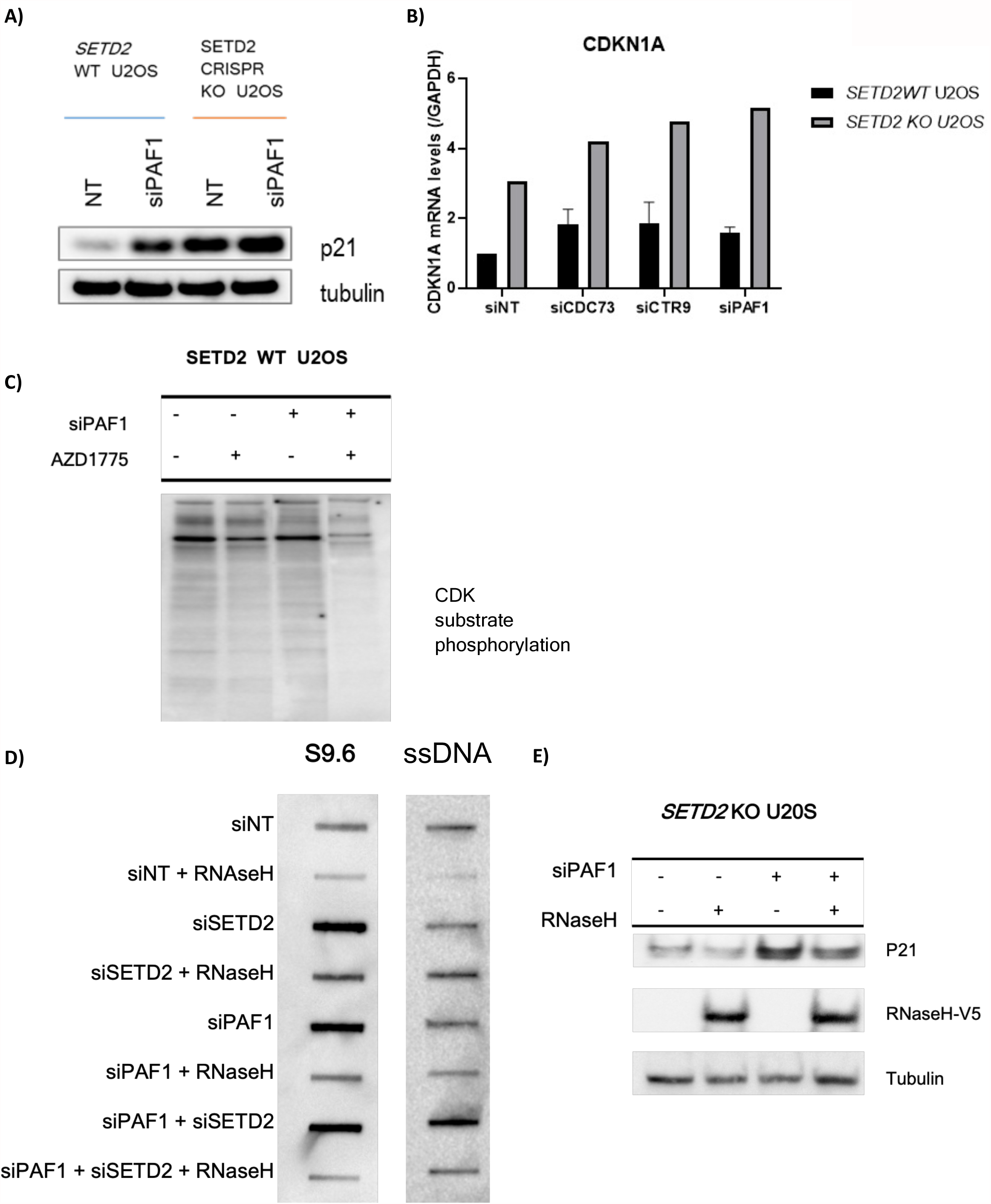
Loss of PAF1 causes R-loop dependent up-regulation of p21 in SETD2-deficient cells. **(A)** Western blot analysis of p21 protein levels in SETD2 WT and SETD2 CRISPR KO U2OS cells after 48 hours of treatment with either non-targeting control siRNA or PAF1 siRNA. Tubulin was used as a loading control. **(B)** qRT-PCR analysis of CDKN1A levels normalised to 18S in U2OS cells transfected with the indicated siRNAs (n = 3). **(C)** Western Blot analysis of p-CDK(S) activity following siPAF1 with and without AZD1775 in *SETD2 WT* U2OS cells. **(D)** RNA:DNA hybrid slot blot of genomic DNA ± RNaseH1 treatment from cells treated with siNT, siSETD2, siPAF1 or combined siSETD2 and siPAF1. **(E)** Western blot analysis of p21 protein levels in SETD2 KO U2OS cells. Immunoblot staining for the V5-epitope confirms RNAseH1 overexpression.

We found that the reduction in long-term colony formation of cells depleted for PAF1 (data not shown) coincided with increased mRNA and protein levels of p21 (Figure 3A-B), a potent inhibitor of Cdk-Cyclin complexes. We do, however, not observe any p21 expression in the absence of PAF1 in *TP53* KO U2OS cells (data not shown). Following loss of PAF1, we found that elevated levels of p21 did indeed reduce CDK activity, as shown by a decrease in phosphorylation of CDK motifs for selected putative substrates, both in the absence and presence of AZD1775 (48 hours post-drug treatment) (Figure 3C). In agreement with this, we find that knockdown of PAF1 restores protein levels of RRM2 following WEE1-inhibitor treatment, consistent with reduced CDK-dependent RRM2 degradation (Supplementary Figure 3C). The above observations suggest that SETD2-CRISPR cells depleted for CDC73, CTR9 or PAF1 exhibit increased p21 levels, reduced CDK1/2 activity and delayed G1-S progression.

Loss of PAF1 has previously been shown to cause an increase in the formation of RNA-DNA hybrids (R-loops) (Shivji *et al*., 2018), which could lead to subsequent checkpoint signalling and p53 activation. Therefore, we asked whether up-regulation of p21, which leads to p53-dependent resistance to WEE1i, could be caused by R-loop formation in the absence of PAF1. First, global R-loop levels were assessed following depletion of SETD2 and/or PAF1 by a slot blot using the anti-DNA:RNA hybrid S9.6 antibody (Figure 3D). This showed an increase in overall R-loop levels in SETD2-, PAF1- and SETD2- and PAF1-depleted U2OS cells. As a control, we pre-treated the genomic DNA with RNase H, which completely abolished the R-loop signal in both control and PAF1-/SETD2-knockdown samples cells (Figure 3D). To test whether the R-loops formed in cells depleted of PAF1 do indeed contribute to DNA damage and p53-mediated activation of p21, we transiently over-expressed V5-RNaseH1, which resolves R-loops, in control- and siPAF1 treated cells (Figure 3E). By analysing p21 levels, we found that expression of RNase H1, as shown by high levels of the V5-epitope tag, resulted in reduction of p21 levels (Figure 3E). Our data suggest that elevated R-loops in the absence of *SETD2* or *PAF1* cause the observed changes in p53-pathway activation, ultimately leading to up-regulation of p21 and G1 arrest.

### Direct role for p21 in regulating sensitivity to WEE1 inhibitor treatment

Next, we sought to determine whether this p21-mediated G1-S delay may be responsible for the observed resistance to WEE1 inhibition in the absence of PAF1. We reasoned that if the G1 delay was mediated by the p53-p21 pathway, then co-depletion of p21 should suppress the cell cycle delay induced by CDC73, CTR9 or PAF1 depletion. We found that depletion of p21 (*CDKN1A*) alone further increased the BrdU^+^ and BrdU^-^ S-phase population (15.2% increase in S-phase compared to siNT-treated *SETD2 KO* U20S cells) following AZD1775 treatment, in accordance with the published role for p21 in regulating the G1/S transition (Abbas and Dutta, 2009) (Supplemental Figure 4A). Furthermore, co-depletion of CDKN1A 24 hours after transfection with PAF1 siRNA completely restored the number of G1 phase cells to control numbers after AZD1775 treatment in SETD2 CRISPR KO U20S cells (Figure 4A, Supplemental Figure 4A). Notably, the low intensity of the BrdU staining in the co-depleted cells (siPAF1 + siCDKN1A), including an increase in the BrdU-negative population, suggested that even though cells depleted for both PAF1 and CDKN1A entered S-phase, replication was still compromised (Figure 4A). These results indicate that p21 is required for the cell cycle effects observed following PAF1 depletion.

**Figure 4.**
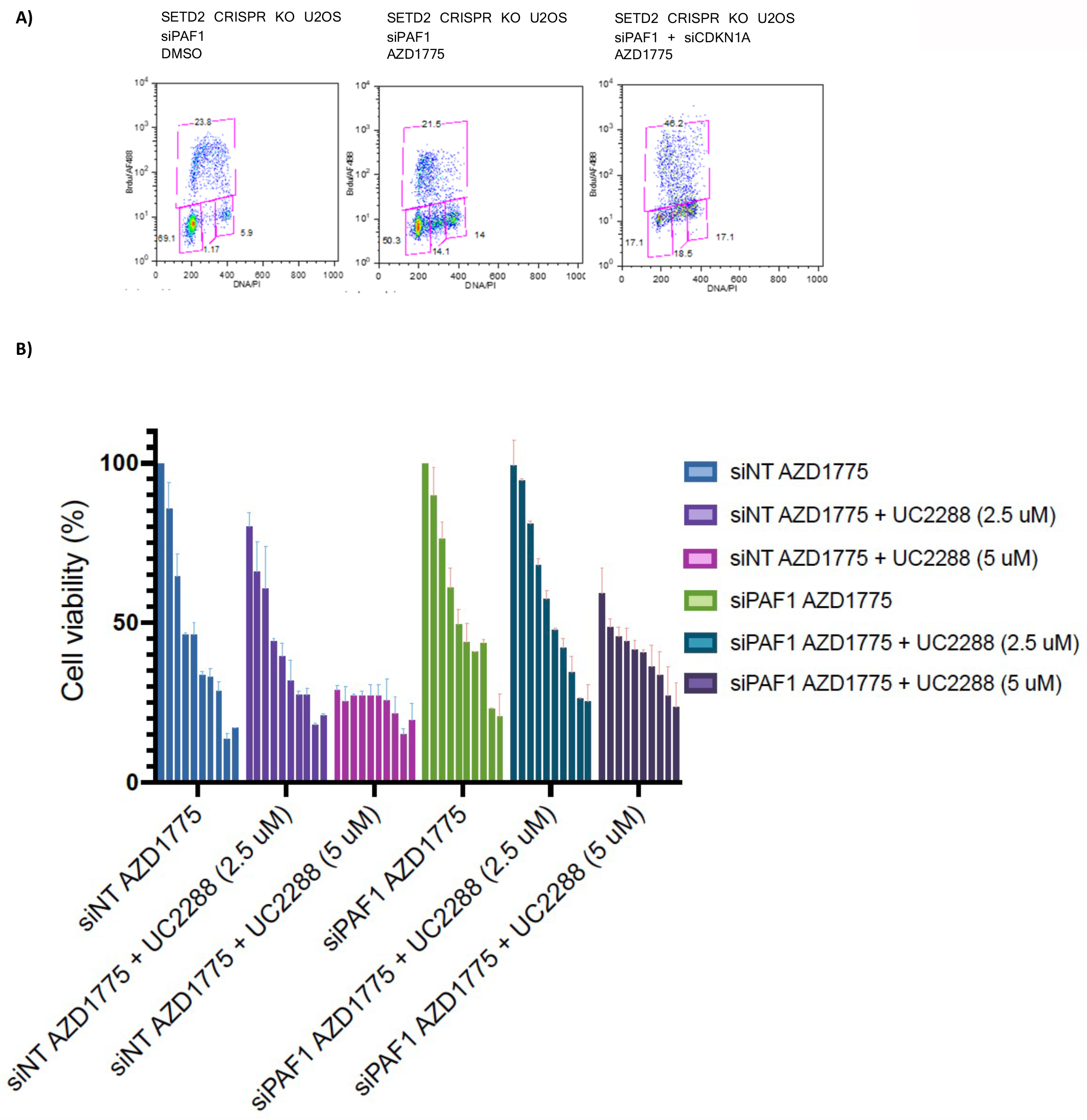
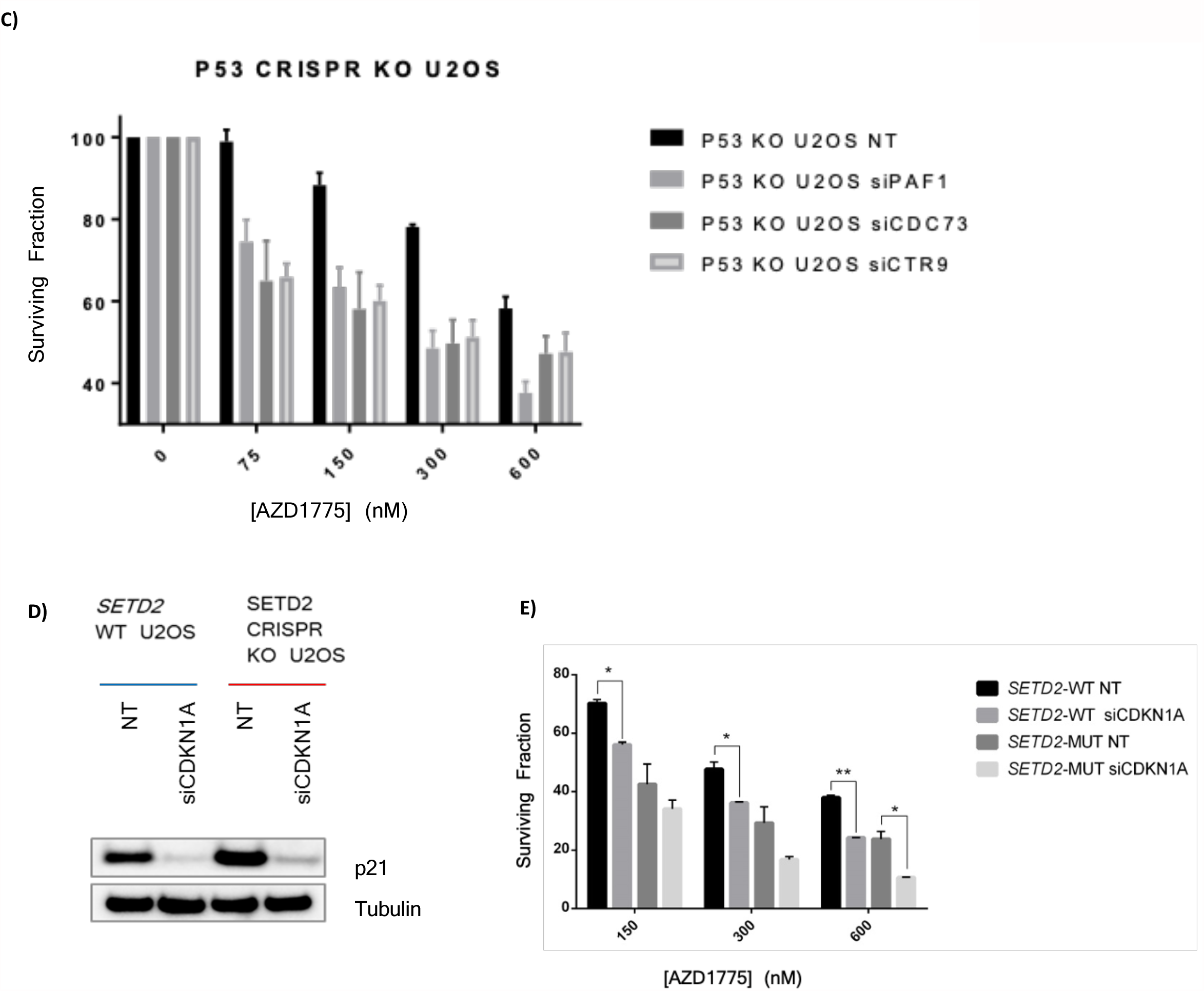
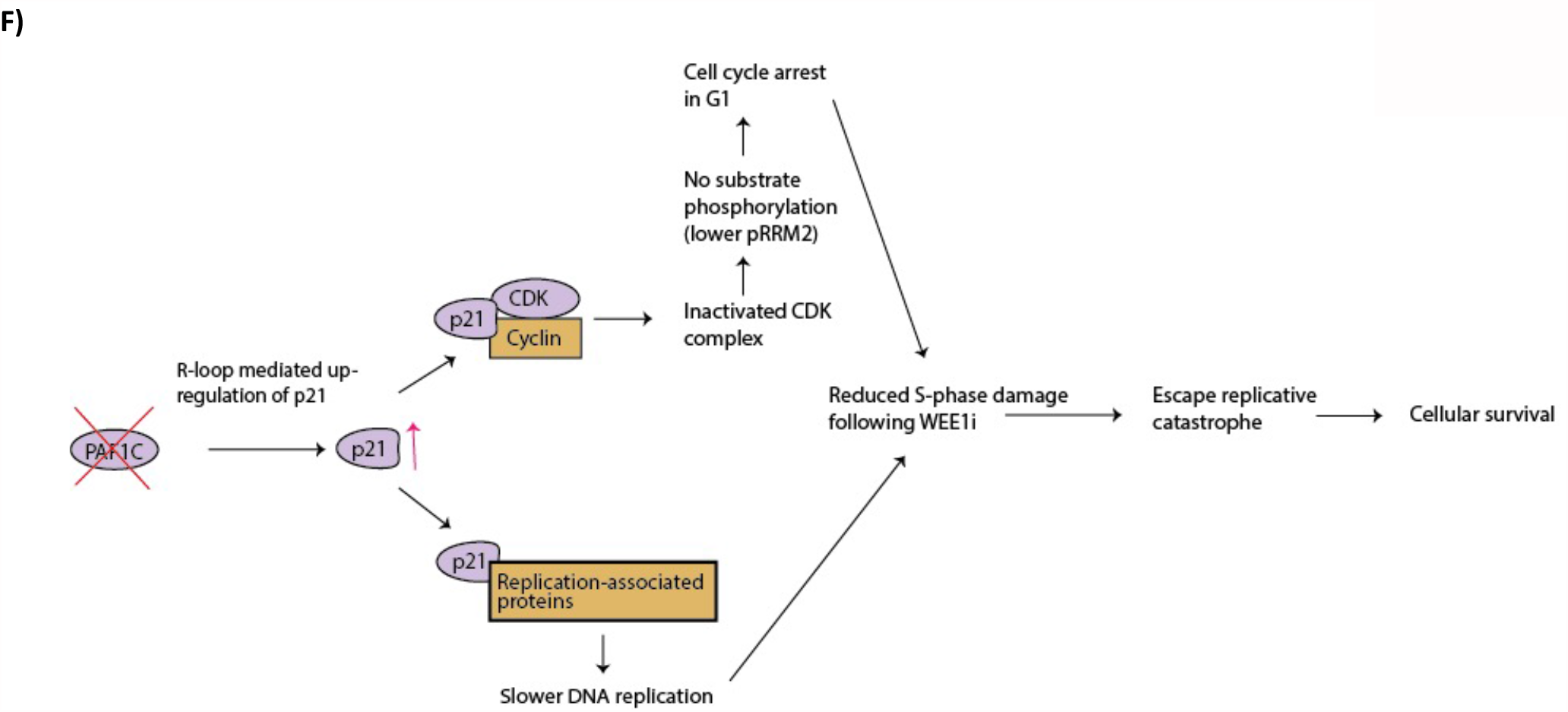
Paf1-mediated suppression is dependent on the p53/p21 axis. **(A)** 48 hr following DMSO -or AZD1775-treatment, siPAF1, or siCDKN1A + siPAF1 treated *SETD2* KO U2OS cells. The data shown are from a single representative experiment out of three repeats. Numbers are the relative percentage of cell cycle stage. **(B)** *SETD2* CRISPR KO U2OS cells treated with siNT or siPAF1 after 48h exposure to different concentrations of AZD1775 (0-10 μM) and UCC2288 (p21 inhibitor). Survival fraction of non-targeting control, CDC73, and CTR9 **(C)** siRNA treated TP53 CRISPR KO cells after 72 h exposure to different concentrations of AZD1775 (75, 150, 300 and 600 nM). Data points and bars represent the mean and SEM of ≥ three independent experiments; * P < 0.05; ** P < 0.01; and *** P < 0.001. **(D)** Western blot analysis of p21 protein levels in SETD2 WT and SETD2 CRISPR KO U2OS cells exposed to either non-targeting control siRNA or p21 siRNA. Tubulin was used as a loading control. **(E)** Cell viability is further reduced when WEE1 inhibition is combined with p21 knock down in SETD2 WT and SETD2 mutant cancer cell lines. **(F)** Model where loss of CDC73, CTR9 and PAF1 results in resistance to WEE1i treatment.

To test whether these p21-induced cell cycle effects were indeed responsible for the PAF1-mediated resistance to AZD1775, we tested whether siPAF1-treated *SETD2* KO U2OS cells could be re-sensitized to WEE1i through inhibition of p21. To this goal, we performed cell cycle analyses and viability assays on *SETD2 CRISPR KO* U2OS transfected with non-targeting control siRNA or PAF1 siRNA using AZD1775 and a pharmacological inhibitor of p21, UC2288, which was shown to downregulate p21 on both the transcriptional and post-transcriptional level (Wettersten *et al*., 2013) (Figure 4B). Indeed, we found that siPAF1-treated cells with high p21 levels could be killed using high concentrations (5 μM) of the p21 inhibitor alone, or with a combination of low doses of AZD1775 and UC2288 (Figure 4B). Next, we reasoned that if loss of PAF1 causes p21 up-regulation in a p53-dependent way, removal of p53 could alleviate p21-mediated G1-arrest in cells depleted of PAF1 components. Therefore, we hypothesized *TP53* KO U2OS cells would display a different response to WEE1-inhibitor treatment when depleted of CDC73, CTR9, or PAF1. Indeed, we found that *TP53* CRISPR KO U2OS cells depleted for CDC73, CTR9 or PAF1 displayed increased sensitivity to WEE1-inhibitor treatment when compared to *TP53* WT U2OS cells (Figure 4C), likely a result of dysfunctional p53 and p21 pathway activation, and therefore further increased CDK activity, leading to increased cell death.

To test whether p21 inhibitors could be applied to specifically target SETD2-deficient tumours, we also explored UC2288 sensitivity in *SETD2* WT versus *SETD2-*mutant RCC cells. First, we measured cell viability in *SETD2* wild type and *SETD2* mutant cells following knockdown of CDKN1A (p21). siRNA treatment significantly reduced p21 protein expression in both cell lines (Figure 4D). As shown in Figure 4E, we found that knockdown of *CDKN1A* further sensitised RCC cell lines 786-0 and A498 to AZD1775 (Figure 4E) In agreement with our findings in RCC cell lines 786-0 and A498, p21 inhibition caused a small increase in the S-phase population (Supplemental Figure 4B) and further sensitised both SETD2 WT and KO U2OS cells to AZD1775, mostly at lower concentrations, showing p21 has a direct role in determining sensitivity to WEE1-inhibitor treatment (Supplemental Figure 4C). Furthermore, we found that *SETD2* KO U2OS cells are more sensitive to p21 inhibition by UC2288 alone when compared to *SETD2* WT U2OS cells, suggesting that p21 is a critical vulnerability in high-p21 expressing SETD2 KO cancer cells.

## Discussion

Here, we characterize CDC73, CTR9, and PAF1, components of the PAF1 complex, as important novel regulators of sensitivity to the WEE1 inhibitor AZD1775. We found that loss of CDC73, CTR9, and PAF1 leads to a partial blockage of the G1/S transition, making cells less prone to replication catastrophe upon treatment with the WEE1 inhibitor. We propose that in the absence of PAF1, CDC73, or CTR9, elevated p21 as a result of high levels of R-loops promotes cell survival through its role as a cyclin-dependent kinase (CDK) inhibitor under replication stress conditions. p21-dependent CDK inhibition results in the partial blockage of G1/S transition thereby slowing of S-phase progression, and ultimately allowing more time for DNA replication and DNA repair. We also show that PAF1-depleted *SETD2*-deficient cells can be re-sensitized to AZD1775 through elevation of CDK activity using a p21 inhibitor.

The effect of CDC73, CTR9, or PAF1 depletion on the replication catastrophe phenotype upon WEE1 inhibition was particularly pronounced in *SETD2-*depleted cells. As elevated CDK activity resulting from Wee1 inactivation is required to manifest the synthetic lethality in between Set2 deletion and Wee1 inactivation in *S. pombe*, we were not surprised to identify loss of function mutations in *cdc2* amongst the spontaneous mutants that suppressed the *set2Δ wee1-50* interaction (Pai *et al*., 2019). CDK1 and CDK2 hyperactivation is central to the manifestation of the synthetic lethality between loss of SETD2 and inhibition of WEE1, as elevated CDK activity promotes increased replication origin firing as well as cyclin F-mediated degradation of RRM2, the regulatory subunit of ribonucleotide reductase (RNR) that catalyses the rate limiting step of the formation of deoxyribonucleotides (D’Angiolella *et al*., 2012; Pfister *et al*., 2015). This ultimately causes cell death due to dNTP shortage. Considering the well-described role of p21 in regulating CDK activity in mammalian cells and its requirement for G1 arrest, we predict that p21 protects cells from AZD1775-induced S phase damage through a similar, but indirect, modulation of CDK activity. P21 also exists in a complex with the proliferating-cell nuclear antigen (PCNA) (Waga *et al*., 1994), providing another way through which p53-dependent induction of p21 can result in slowing of DNA replication.

Previous studies have shown that ATR inhibition could be rescued with a CDK1 inhibitor that prolonged the cell cycle, thereby providing more time for DNA replication such that mitosis and cell division could complete with more fully replicated DNA (Eykelenboom *et al*., 2013). Interestingly, though, recent work by Hauge *et al*., showed that high p21 did not alter S-phase CDK activity but rather the protective effect of p21 in the presence of AZD1775 was mediated through damage reduction, as measured by γH2AX (Hauge, Macurek and Syljuåsen, 2019). They hypothesised that p21 is involved in restraining replication after WEE1 inhibition, limiting S-phase damage and subsequent cell death. Indeed, this hypothesis aligns with previous work from Gottifredi *et al*., where it was shown that only cells expressing low p21 immediately progress through the cell cycle upon release from S phase arrest, whereas high p21 cells move much more slowly through the cell cycle, demonstrating that p21 is required for efficient replication restart (Gottifredi *et al*., 2004)

The direct link between loss of components of the PAF1C and elevated levels of p21 is not yet fully understood, but we propose that high p21 levels following PAF1 depletion are a result of increased R-loop formation. PAF1 loss results in reduced γH2AX and ATM/DNAPK phosphorylation levels and reduced replicative catastrophe in the presence of AZD1775 at 48 hours, suggesting delayed or reduced DNA double-strand break formation, probably as a result of more complete DNA replication. We do, however, observe p53-dependent CDKN1A up-regulation between 24 and 48 hours post-AZD1775 treatment in the absence of PAF1, perhaps in an ATR- and Chk1-dependent manner. Therefore, we propose that elevated R-loops in PAF1-depleted cells lead to replication fork slowing (reviewed in Gómez-González & Aguilera, 2019), alleviating the lethal effects of replication stress in the first S-phase, whereas under-replicated DNA and unrepaired damage, possibly as a result of elevated transcription-replication conflicts, are carried over into the next cell cycle, leading to problems in mitosis and elevated p21 levels in the subsequent G1 phase.

We therefore propose the following model, where loss of components of the PAF1 complex, including CDC73, CTR9 and PAF1, result in up-regulation of p21 through direct PAF1-mediated up-regulation of R-loop formation. High p21 levels are subsequently responsible for 1) slowing down of replication initiation and progression (Hauge et al., 2019; Waga, Hannon, Beach, & Stillman, 1994) and 2) reduction of CDK activity, leading to a second-cycle G1 arrest, both of which ultimately lead to reduced S-phase damage, rescue of replication catastrophe and increased cellular survival in the presence of AZD1775 (Figure 4F).

Even though PAF1-depleted cells display cross-resistance to other inhibitors of ribonucleotide reductase, including HU and gemcitabine, they showed reduced cellular survival upon γ-irradiation, suggesting a complex relationship between the PAF1 complex, nucleotide pools and the response to DNA damage. High levels of p21 in PAF1-depleted cells might, in addition to causing prolonged G1 arrest in the subsequent cell cycle, also suppress the degradation of RRM2 via inhibition of the CDK2-cyclin E complex, thereby restoring nucleotide pools. In *S. pombe*, previous studies have found that high dNTP levels can lead to increased DNA damage sensitivity and slow progression through S-phase (Fleck *et al*., 2013; Pai and Kearsey, 2017). A recent study has suggested a role for PAF1 in attenuating radiosensitivity by inhibiting immediate-early response 5 (IER5) transcription, which is involved in cell cycle regulation (Ding *et al*., 2019; Zheng *et al*., 2020).

Our results suggest that tumours with high levels of R-loops and concomitant increases in p21 caused by low levels of CDC73, PAF1, and CTR9 would not benefit from treatment with the WEE1 inhibitor. Previous studies have implicated mutations in members of the PAF1C, including CTR9, in tumorigenesis (Hanks *et al*., 2014). As AZD1775 is currently being investigated in more than fifty clinical trials for cancer treatment, as monotherapy or in combination with chemotherapeutic drugs and/or radiation (www.clinicaltrials.gov), expression levels and mutation status of PAF1C components should be taken into consideration in the clinical context. Finally, we propose that a combination of low doses of AZD1775 (WEE1i) and UC2288 (p21i) could be used to improve the therapeutic response of SETD2-deficient cells, even in the context of PAF1 loss, which should be further explored in *in vivo* studies.

## Materials and Methods

### Yeast strains and media

Strains were cultured and stored as described in Moreno, Klar, & Nurse, 1991. YE6S medium contained 3% glucose, 0.5% EZmixTM yeast extract (Sigma), and 1.125 g/L supplements mix containing equal amounts of adenine, arginine, L-histidine, L-leucine, lysine, tryptophan, and uracil. Strains used in this study can be found in Table 1.

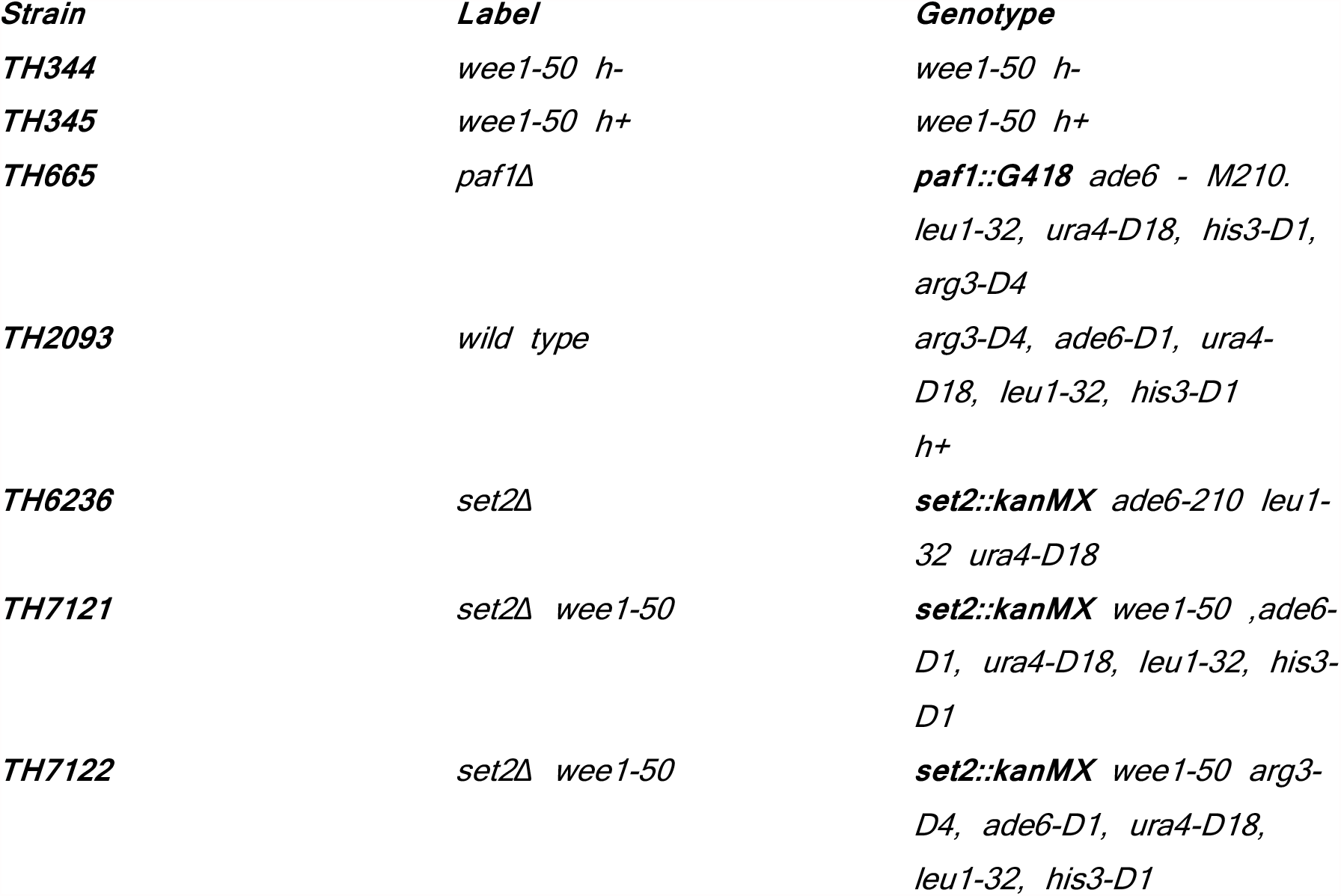

### Serial Dilution Assay

A dilution series for the indicated mutant cells was spotted onto YES plates. Plates were incubated at 25°C, 32°C, or 36°C for 2–3 days before analysis.

### Cell culture

U2OS (human osteosarcoma) and the human glioblastoma cell line T98G cells were obtained from ATCC (ATCC number HTB-96 and CRL-1690). The U2OS *SETD2* CRISPR knockout (KO) cell line was generated using the CRISPR-Cas9 technology as previously described (Pfister et al., 2015). The target sequence of the chimeric gRNA scaffold was ACTCTGATCGTCGCTACCAT (first exon of *SETD2*) (Pfister et al., 2015). The U2OS P53 CRISPR KO cell line was obtained from Dr. Kamila Burdova (V. D’Angiolella Lab). Human renal cell carcinoma cell lines 786-O (*SETD2* wild-type) were a kind gift from V. Macaulay and A498 (SETD2 homozygous truncating mutation) cells were obtained from ATCC (ATCC number HTB-44). Human cancer cell lines were cultured in Dulbecco’s Modified Eagle’s Medium (DMEM) supplemented with 10% v/v foetal bovine serum (FBS, Sigma), penicillin (100 units/mL, Sigma) and streptomycin (0.1 mg/mL, Sigma) at 37°C in a humidified atmosphere containing 5% CO2.

### Transient siRNA-mediated gene knockdown

Cell lines were reverse transfected with short interfering RNAs (siRNAs) (12 nM final concentration) using Lipofectamine RNAiMAX (Invitrogen) according to the manufacturer’s instructions. Medium was replaced 16-20 hours after transfection. The target sequences of the siRNAs used in this project are listed below: siCDC73 (ON-TARGET plus SMARTpool, 015184), siCDKN1A (ON-TARGET plus SMARTpool, 003471), siCTR9 (ON-TARGET plus SMARTpool, 032246), siLEO1 (ON-TARGET plus SMARTpool, 016579), siPAF1 (ON-TARGET plus SMARTpool, 020349), siRTF1 (ON-TARGET plus SMARTpool, 014104), and siSETD2#3 (si#3) (Dharmacon): GAAACCGUCUCCAGUCUGU, and siSETD2#5 (si#5) (Dharmacon): UAAAGGAGGUAUAUCGAAU.

### Drug treatment

All inhibitors were dissolved in Dimethyl sulfoxide (DMSO) and stored at - 80 °C per the manufacturer’s instructions. Inhibitors used in this study are AZD1775 (MK-1775) (Axon Medchem) and Gemcitabine (Sigma Aldrich).

### Clonogenic Survival Assays

Cells were serially diluted into 6-well plates and allowed to attach for 24 hours. They were then exposed to ionising radiation (IR) and grown in fresh medium for 10 days. Colonies were stained with crystal violet and manually counted. Clonogenic survival was expressed as % colonies relative to untreated cells.

### Quantitative RT-PCR (Reverse Transcription-Polymerase Chain Reaction)

Total RNA was purified from cell pellets with the RNeasy mini kit (Qiagen). RNA concentration was determined using a Nanodrop 1000 spectrophotometer (Thermo Scientific). Purified total RNA (1 μg) was reverse transcribed using the superscript iii first strand synthesis supermix for qRT PCR kit (Invitrogen), according to the manufacturer’s instructions. Quantitative RT-PCR was performed using Absolute Blue QPCR SYBR green, low ROX Mix (Thermo Scientific). The comparative CT method was applied for quantification of gene expression. Primers are available upon request.

### Resazurin assay for cell viability

48 hours following drug treatment, culture medium was removed and fresh medium containing 10 μg/mL resazurin was added to each well. The fluorescent signal was measured by a fluorescence plate reader (BMG Labtech POLARstar OMEGA) after 2 hours of incubation at 37 °C.

### Fluorescence-activated cell sorting (FACS) analysis

Cells were collected by trypsinization and fixed in 70% ice-cold ethanol (EtOH). Fixed cells were resuspended in 1 mL PBS supplemented with 10 μL 1 mg/mL propodium iodide (PI). Cells were incubated for >30 min at RT and then analysed for DNA content by flow cytometry. For simultaneous staining of gH2AX and EdU, cells were labelled with gH2AX as and EdU as previously described (HB *et al*., 2020). Briefly, cells were incubated for 1 h with 2 μM EdU (Thermo Fisher) and fixed in 70% ethanol, prior to labelling with primary (gH2AX antibodies (clone JBW301, Millipore)) and secondary (anti-mouse Alexa488, A21202 life technologies) antibodies. EdU was detected using the Click-iT Plus EdU Alexa Fluor 594 Flow Cytometry Assay Kit (Thermo Fisher). For the experiments in Figure 2C-D, a barcoded (with Alexa Fluor 647) control sample of non-treated cells was added to all the individual samples, to eliminate sample to sample variation during staining. Samples were analysed in a LSRII flow cytometer (BD Biosciences) and processed in FACSDiva and FlowJo software (Both BD Biosciences).

### Immunoblotting

Cells were lysed in lysis buffer (50 mM Tris HCl pH 7.5, 150 mM NaCl, 1% Triton) supplemented with 1:50 fresh protease inhibitor cocktail and 1:100 fresh phosphatase inhibitor cocktail (1mM NaOVa; 10mM NaF; 10mM NaPyrophosphate) and immunoblotting was performed as described previously. Targets of the primary antibodies used for western blotting were as follows: Apoptosis Cocktail (pro/p17-caspase-3, cleaved caspase-3, cleaved-PARP, cleaved-actin, 1:500, Abcam), RRM2, p21 (1:1000, Cell Signalling), p-CDK(S), p-ATR(S428), ATR, p-Chk1(S317), Chk1, P-ATM S1981 (Cell signalling). DNA-PK pS2056 (abcam), p-RPA S4S8 (Bethyl), CDK1 (sc-54 Santa Cruz biotech), pCDK1 Y15 (Cell signalling), PAF1 (Cell Signaling), Lamin B (cell signalling), GAPDH and Tubulin. Stain free technology (Biorad) was used for total protein detection in Figure 2E.

### Slot Blot

U2OS cells were lysed in 200 uL lysis buffer (100 mM Tris-HCl [pH 8.5], 5 mM EDTA, 0.2% SDS, and 100 mM NaCl) containing 0.5 mg/ml proteinase K and left at 55°C, 350 RPM O/N. Genomic DNA was precipitated with isopropanol for 5 min RT, spun down for 3 min at 3000g (4°C,) washed with 500 µL 70% ethanol and spun down at for 3 min at 3000g (4°C,). Supernatant was removed and samples were air-dried. Pellets were resuspended in 100 - 200 uL Tris-EDTA (TE) buffer. gDNA was mixed well and left at 55°C for 10 min to dissolve. All samples were diluted to give 50 µg gDNA in a volume of 100 µL TE. 2uL RNase A (10 mg/mL, Thermo Cat# EN0531) was added to each and incubated 2 hr 37°C, 300 RPM. For the RNaseH1 control, 20 µL gDNA was added to 25 µL TE + 5uL of 10X RNaseH1 buffer + 2uL RNaseH1 and incubated O/N 37°C, 300 RPM. The remaining untreated gDNA was also incubated O/N 37°C, 300 RPM. The gDNA samples (5 µg for S9.6 blot and 0.5 µg for ssDNA blot) were blotted onto a positively charged nylon transfer membrane (GE Healthcare Cat# RPN203B) using a slot blot apparatus (Bio-Rad). ssDNA blot was incubated in denaturing buffer (0.4 M NaOH and 0.6 M NaCl), shaking for 10 min RT, followed by neutralizing buffer (1.5 M NaCl and 0.5 M Tris [pH 7.4]) shaking for 10 min RT. Blots were quickly dried using wattman paper and UV crosslinked using (0.12 J/m2). Blots were probed O/N using S9.6 (1:1000) or ssDNA-specific antibodies (1:5000), washed in TBS-T, secondary antibody (Anti-Ms-HRP, 1:5000, 2hr RT) and imaged.

### Statistical analysis

Unless stated otherwise, all data in the figures and corresponding text are expressed as the mean +/-SEM of n samples (n ≥ 3). Student’s unpaired t-tests were used for comparisons between two or more experimental groups, without assuming a consistent SD. P-values of <0.05 were considered statistically significant.

## Figure Legends

**Supplemental Figure 1.**
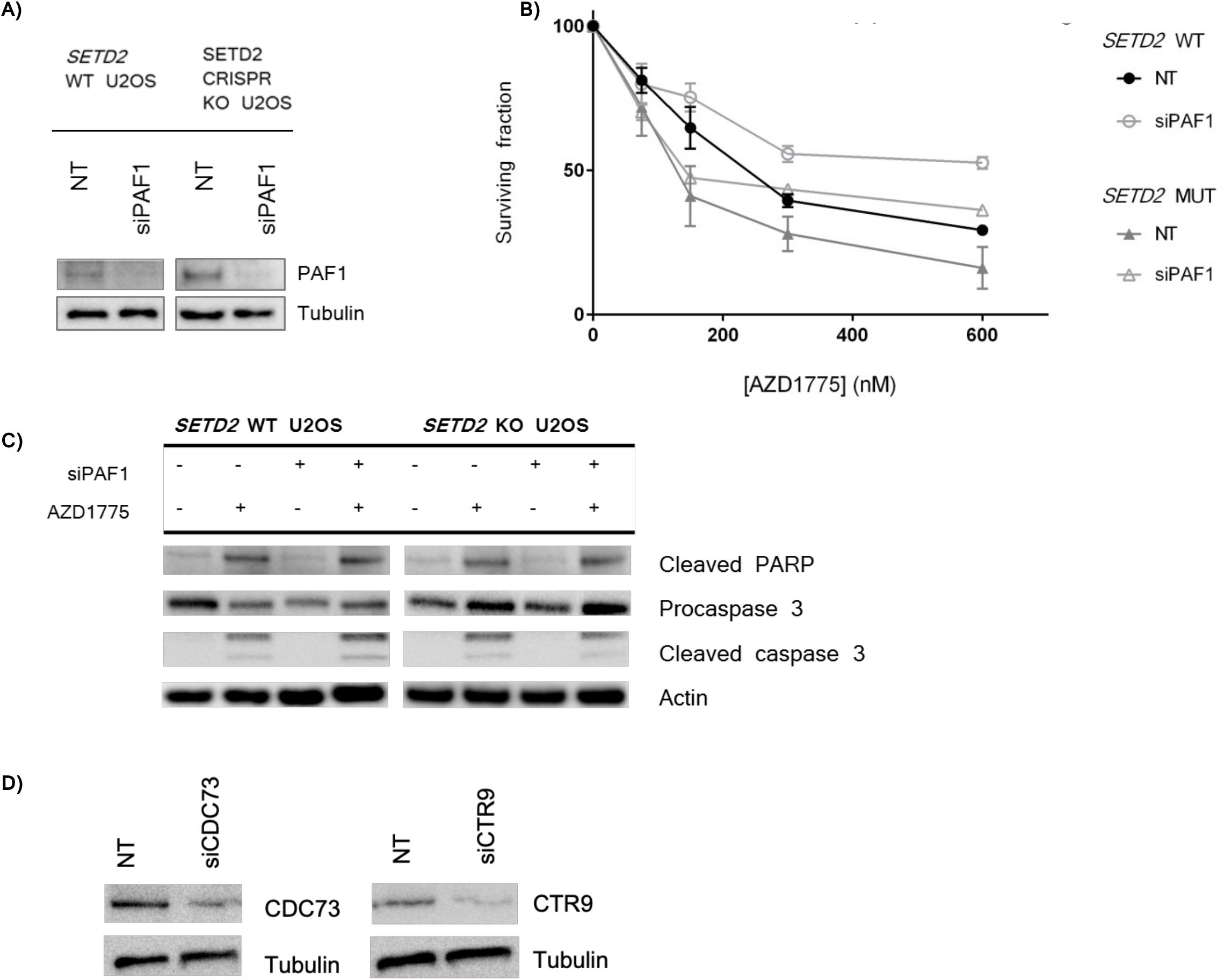
**(A)** Validation of PAF1 siRNA knockdown efficiency by Western Blot in SETD2 WT and SETD2 CRISPR KO U2OS cells 72 hours post-siRNA treatment. **(B)** Viability assay to examine the impact of PAF1 silencing on AZD1775-sensitivity. Survival fraction of non-targeting control and PAF1 siRNA treated (A-B) 786-0 (*SETD2* WT) and A498 (SETD2-mutant) renal cancer cells. **C)** Cleaved PARP, procaspase 3, cleaved caspase 3 and actin protein expression in SETD2 WT and SETD2 CRISPR KO U2OS cells 48 hours post-treatment with either 200 nM DMSO or 200 nM AZD1775 was assessed using Western Blot. **D)** Validation of CDC73 and CTR9 siRNA knockdown efficiency by Western Blot in SETD2 WT and SETD2 CRISPR KO U2OS cells 72 hours post-siRNA treatment.

**Supplemental Figure 2.**
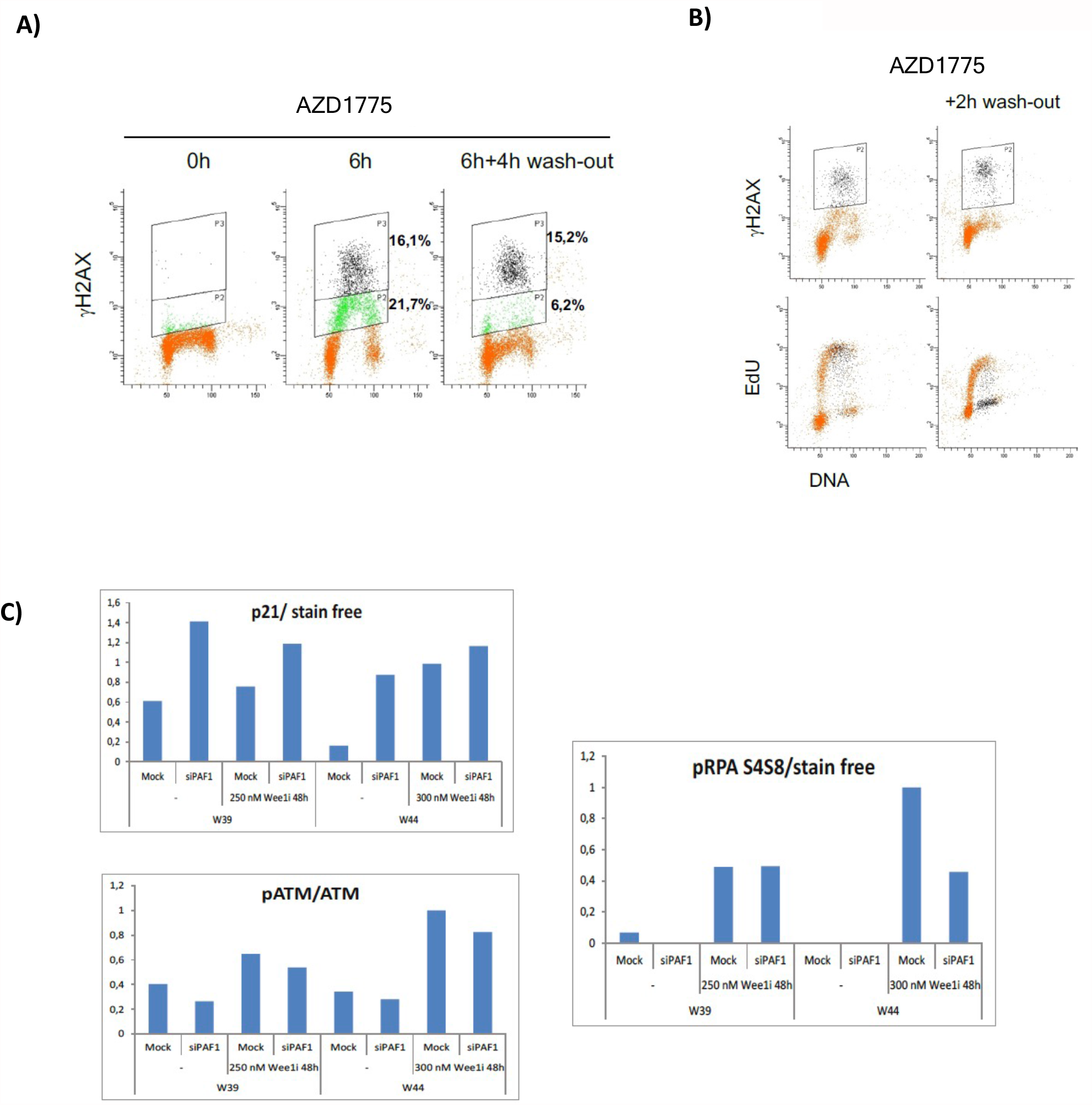
**(A)** yH2AX level measured by flow cytometry after wash-out of MK-1775. U20S cells were treated with 1 µM MK-1775 for 6h, then washed with PBS and cultured further for 4 h. Cells were fixed and stained for yH2AX and DNA content. The strong and intermediate yH2AX populations were gated and percentages determined. One representative experiment out of 15 experiments using different MK-1775 concentration. **(B)** The strong yH2AX population ceases replication. U20S cells were treated with 1 µM MK-177 5 for 7h, washed in PBS and cultured further for 2h. For the last hour 1 µM EdU was added. Cells were fixed and stained for yH2AX, EdU and DNA content. The strong yH2AX population was gated and is indicated in black in the EdU plot. **(C)** Quantification of p21, p-ATM/ATM and pRPA levels from Western Blot shown in Figure 2E.

**Supplemental Figure 3.**
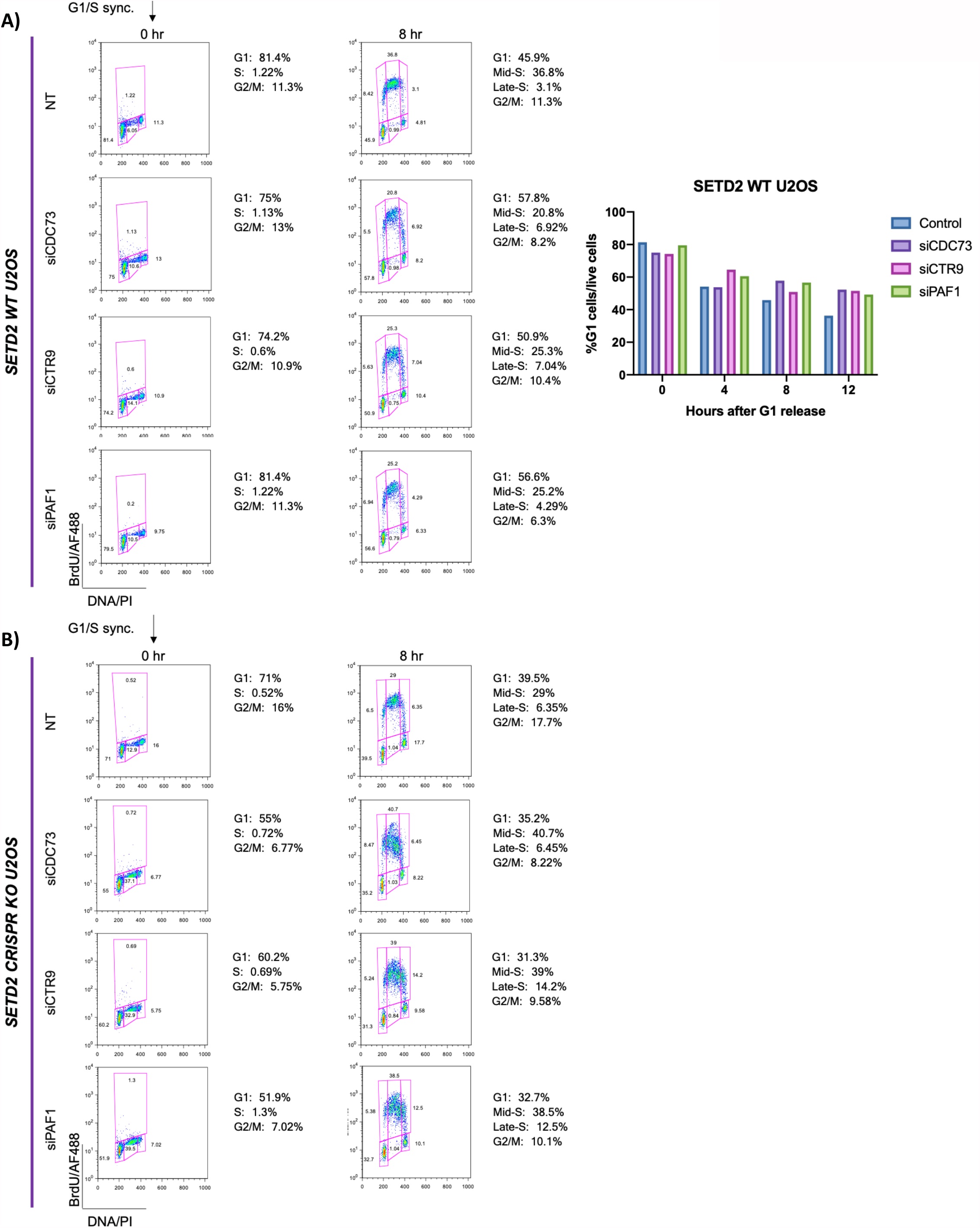
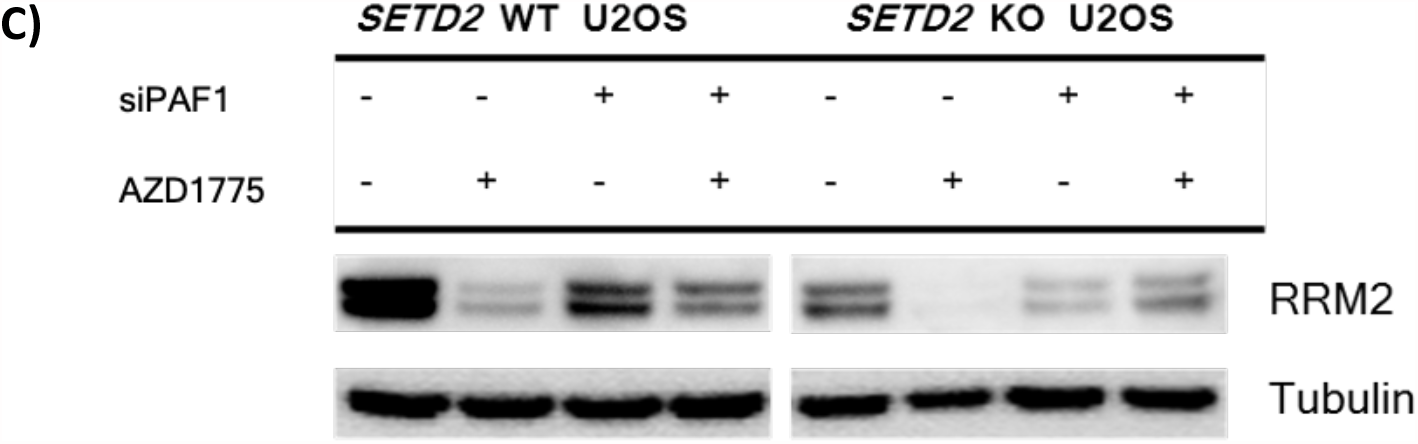
**(A)** *SETD2* WT and **(B)** *SETD2 CRISPR KO* U2OS were treated with control, siCDC73, siCTR9 or siPAF1 and subsequently synchronized at the G_1_/S transition by a double thymidine block (DTB) and then released by addition of fresh medium. Cells were incubated with BrdU for 30 min and collected for cell cycle analysis 4, 8, and 12 hours after release. **(C)** Western blot analysis of RRM2 protein levels in *SETD2* WT and *SETD2* CRISPR KO U2OS cells exposed to either non-targeting control siRNA or PAF1 siRNA and/or treated with either DMSO or 300 nM AZD1775. Tubulin was used as a loading control.

**Supplemental Figure 4.**
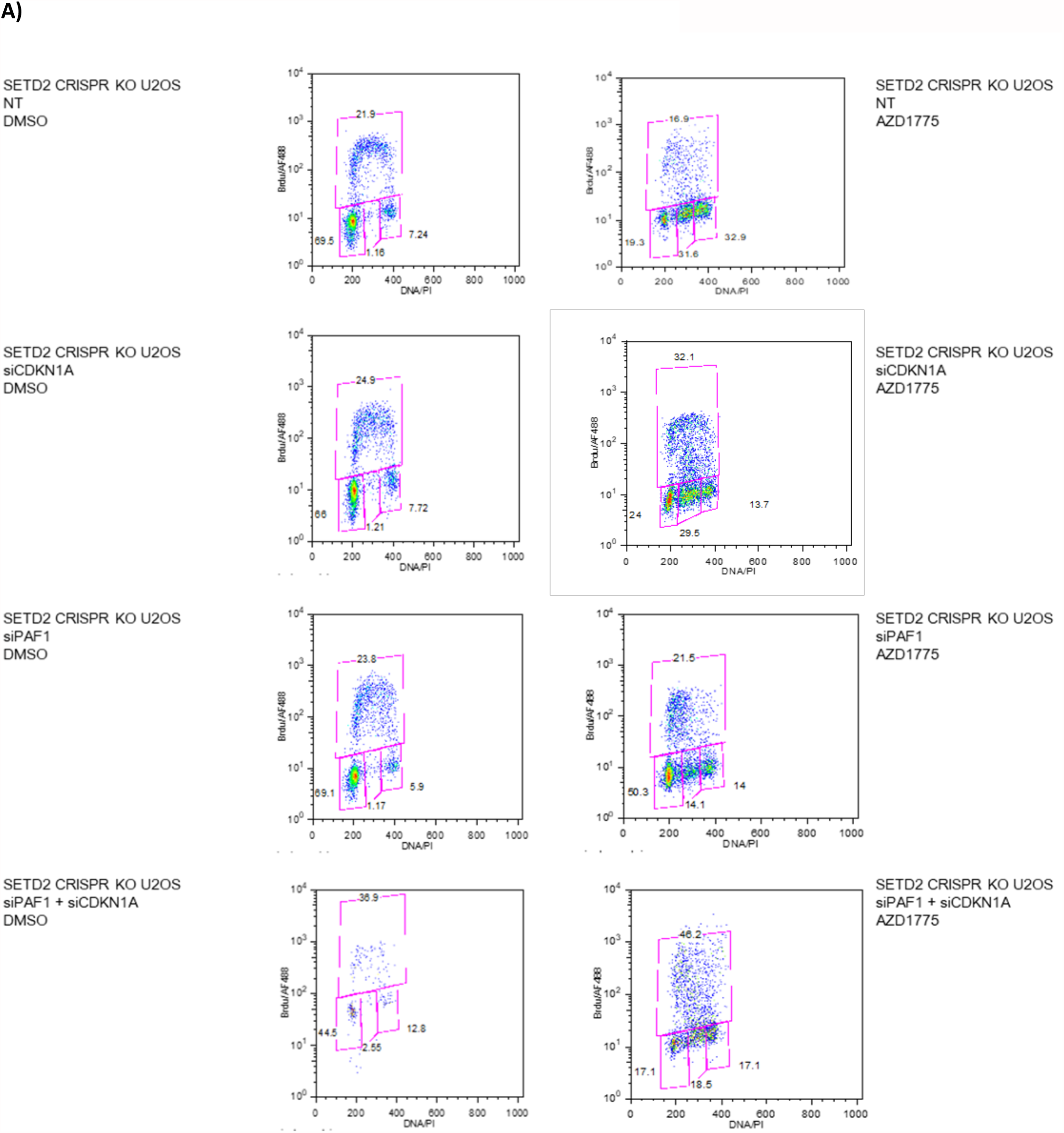
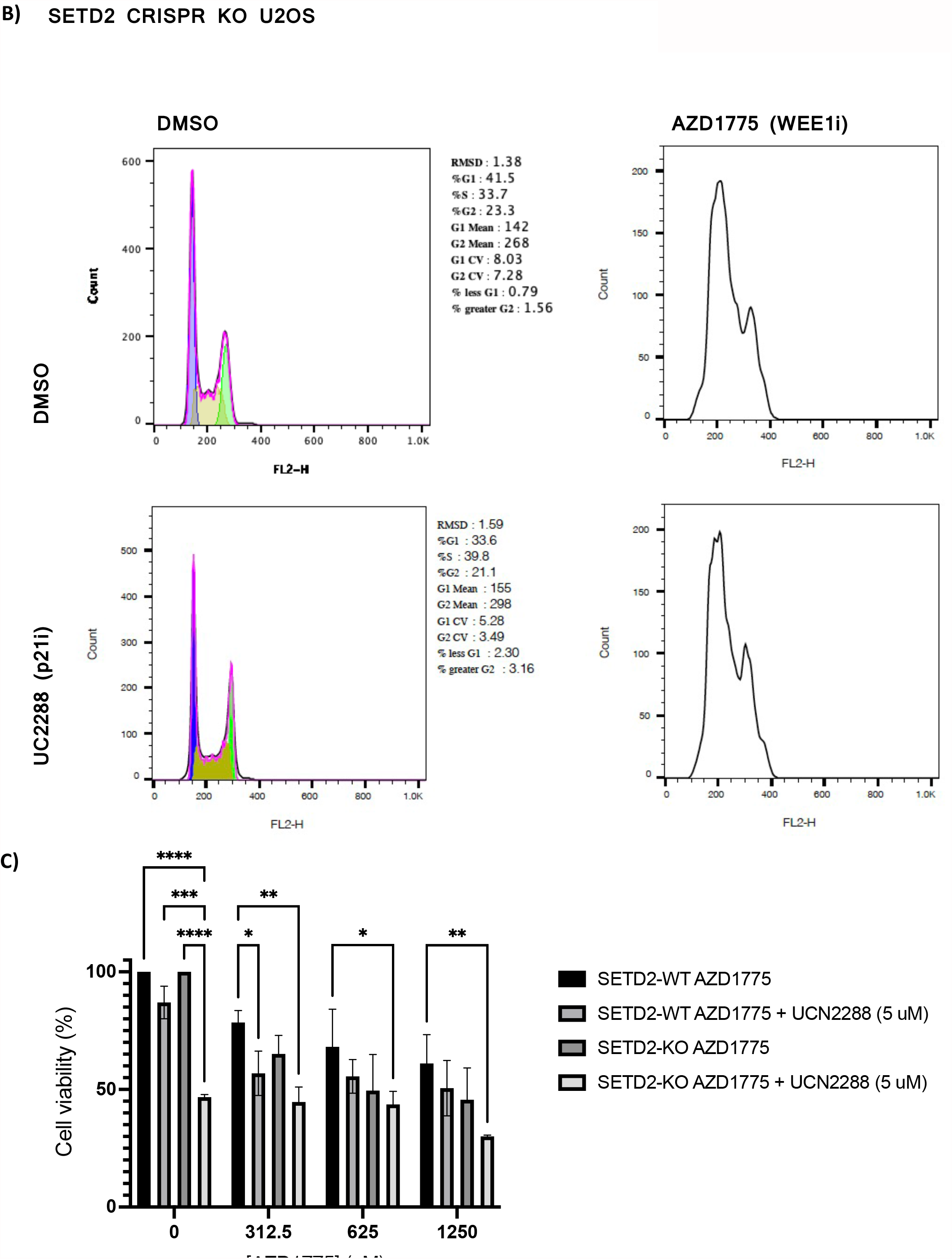
**A)** 48 hr following DMSO -or AZD1775-treatment, control, siCDKN1A, siPAF1, or siCDKN1A + siPAF1 treated *SETD2* KO U2OS cells were pulse-labelled with BrdU for 30 min and collected for cell cycle analysis. The data shown are from a single representative experiment out of three repeats. Numbers are the relative percentage of cell cycle stage. **B)** 48 hr following DMSO, AZD1775- and/or UC2288 treatment, *SETD2* KO U2OS cells were collected for cell cycle analysis. **C)** *SETD2* WT and *SETD2* CRISPR KO U2OS cells after 48h exposure to different concentrations of AZD1775 (0-10 **μ**M) and UCC2288 (p21 inhibitor).

